# Characterizing the transcriptomic response to interferon and infection in European Domestic Ferret respiratory tissues using long-read RNA sequencing

**DOI:** 10.1101/2025.07.21.666063

**Authors:** Rubaiyea Farrukee, Jessie J.-Y. Chang, Jianshu Zhang, James Barnes, Shu Xin Zhang, Sher Maine Tan, Patrick Reading, Lachlan J. M. Coin

**Author notes:** Corresponding authors/Lead contacts. These authors contributed equally (Rubaiyea Farrukee, Jessie J.-Y. Chang, Jianshu Zhang).

## Abstract

The European domestic ferret (*Mustela putorius furo*) is considered the gold-standard small animal model for studying human and avian influenza virus infections. However, experimental characterization of the transcriptomic response to interferon (IFN) stimulation and/or influenza virus infection has been limited, particularly in defining the induction of interferon-stimulated genes (ISGs), with most being computationally predicted. In this study, we present a comprehensive transcriptome-wide assessment of the ferret transcriptome following IFN-α treatment of a ferret lung (FRL) cell line, as well as in nasal turbinates from influenza A virus (IAV)-infected ferrets using long-read RNA sequencing. We have identified a panel of ferret genes orthologous to human ISGs that are upregulated both in response to IFN-α stimulation *in vitro* and IAV infection *in vivo*. We have also identified novel interferon stimulated genes and transcripts. Furthermore, we observed elongation of the poly(A) tails of genes in the *ribosome* and *Coronavirus Disease-19* pathways in response to IFN-α treatment *in vitro,* suggesting a relationship between poly(A) elongation and the antiviral responses of the host. These results illuminate the dynamics of the transcriptional innate immune response of the domestic ferret and provide an important resource for better utilizing ferrets as a small animal model to study influenza virus infections.

## Introduction

Ferrets are widely regarded as the gold-standard animal model for studying influenza virus infections *in vivo* and offer a number of advantages over conventional mouse models (1). Ferrets share similarities with humans in their lung architecture and in the distribution of particular sialic acid receptors in their airways and can therefore be infected with a wide range of human influenza A viruses (IAVs) without the need for prior host adaptation (1,2). Also, unlike mice, ferrets show clinical signs of infection (sneezing, fever, lethargy etc.) and can transmit virus between animals via direct contact or by aerosols, making them an ideal model for measuring impact of therapeutic interventions on clinical outcomes of IAV infection and transmission. In addition, ferrets are outbred animals that show a high degree of genetic similarity (>99% at mitochondrial level) to European minks (*Mustela luterola*), which represent an intermediate reservoir species for avian IAV infections (3). This was highlighted in the 2022 outbreak of highly pathogenic avian influenza (HPAI) H5N1 virus (clade 2.3.4.4b) in mink farms in Spain which resulted in the culling of >50,000 animals (4). While previous studies confirm that mink are susceptible to both human and avian IAV (5), few experimental studies have used mink whereas ferrets are widely used and therefore serve as an ideal surrogate model for mink.

Despite the well-established use of ferrets to study pathogenesis and immunity to influenza virus, limited tools and reagents are currently available to characterise the immune responses elicited in these animals. Studies examining innate immune responses to virus infection, characterized by the secretion of type I and III interferons (IFNs) and upregulation of interferon-stimulated genes (ISGs), are limited and generally rely on qPCR to detect gene expression (6,7). In humans and mice, many ISGs are known to mediate antiviral activity against a broad range of viruses, including against IAV, however less is known regarding their role in other species such as ferrets. We recently characterized ferret Mx1 (an ISG which shows 100% sequence identity to European mink Mx1 at the mRNA level), reporting that it differed significantly to human Mx1 in the potency of its antiviral activity against human seasonal IAV (8). These findings highlight key differences in the function of a particular ISG between humans and ferrets/mink. However, the ferret genome and the transcriptome are poorly annotated on National Center for Biotechnology Information (NCBI) database, making it difficult to identify ferret orthologous of additional human ISGs with known antiviral activity against IAV (8). It remains important to characterize ferret orthologues to define similarities and differences between human and ferret ISGs as such studies have important implications for contextualizing the relevance of conclusions drawn from ferret studies to human influenza infections. Improved annotation of the ferret genome and transcriptome, as well as functional validation of key ISGs, would also enhance the utility of the ferret model for evaluating the efficacy of antiviral therapies and vaccines targeting innate immune pathways (9). Moreover, a better understanding of the ferret ISG repertoire may also reveal host-specific ISG responses that can shape viral evolutionary trajectories. This knowledge is especially relevant for zoonotic viruses such as avian IAVs, which may undergo adaptive changes in intermediate hosts such as ferrets and mink before acquiring the capacity to infect humans more efficiently. Thus, bridging the current knowledge gap in ferret innate immunity is essential to maximise the utility of this valuable animal model and to advance our understanding of host adaptation of influenza viruses.

Third-generation sequencing platforms such as Oxford Nanopore Technologies (ONT) allow the full-length sequencing of RNA transcripts, enabling an enhanced view of the whole transcriptome (10,11). The technology is particularly useful for detecting novel transcripts due to accurate elucidation of all possible alternative splicing events on one single strand of RNA, which is arduous and near impossible using short-read sequencing platforms such as Illumina (12). Therefore, as most of the ferret transcriptome is still largely predicted, ONT long-read sequencing provides an efficient method to validate long-transcripts with multiple alternative splicing events. This ability extends to visualizing trans-spliced transcripts, which are transcripts derived from two pre-mRNA molecules, during the splicing process (13). Furthermore, an additional advantage of ONT RNA-sequencing is its ability to capture the full length of the poly(A) tail on transcripts (12–14), which is also limited in short-read sequencing (17). Poly(A) tails on mRNA are known to be associated with various molecular functions, such as mRNA circularization, localization, decay rates, stability, and regulation of gene expression (14,16,18–22). While short-read sequencing has been previously used to define the ferret transcriptome (23–25) and the diversity of its B cell receptors (26), to our knowledge, a thorough investigation into the transcriptome and polyadenylome of domestic ferret following IAV infection using long-read sequencing has not been reported. Thus, in this study, we sought to utilize the advantages of long-read Nanopore sequencing to comprehensively interrogate the transcriptome of the European domestic ferret (*Mustela putorius furo*).

## Results

### General features and metrics of the domestic ferret transcriptome

To characterise the ferret transcriptome, samples were derived from ferret lung epithelial (FRL) cells cultured for 24 hr in the presence or absence of recombinant ferret IFNα, as well as nasal turbinates isolated from ferrets 6 or 24 hours post infection (hpi) after intranasal infection with PBS (mock) or IAV strain A/Perth/269/2009 (Perth/09, H1N1pdm09) (**Figure 1A**). Following RNA extraction, samples were sequenced using the ONT PCR-cDNA Barcoding kit. As the ferret transcriptome is still largely incomplete, we investigated the presence of hypothetical (‘XM’) transcripts (as annotated on NCBI), and unannotated novel transcripts. A total of ∼46.3 M passed reads were sequenced and re-stranded, with a N50 read length of 743 nt and median length quality score of 12.6 (**Figure 1B**).

**Figure 1.**
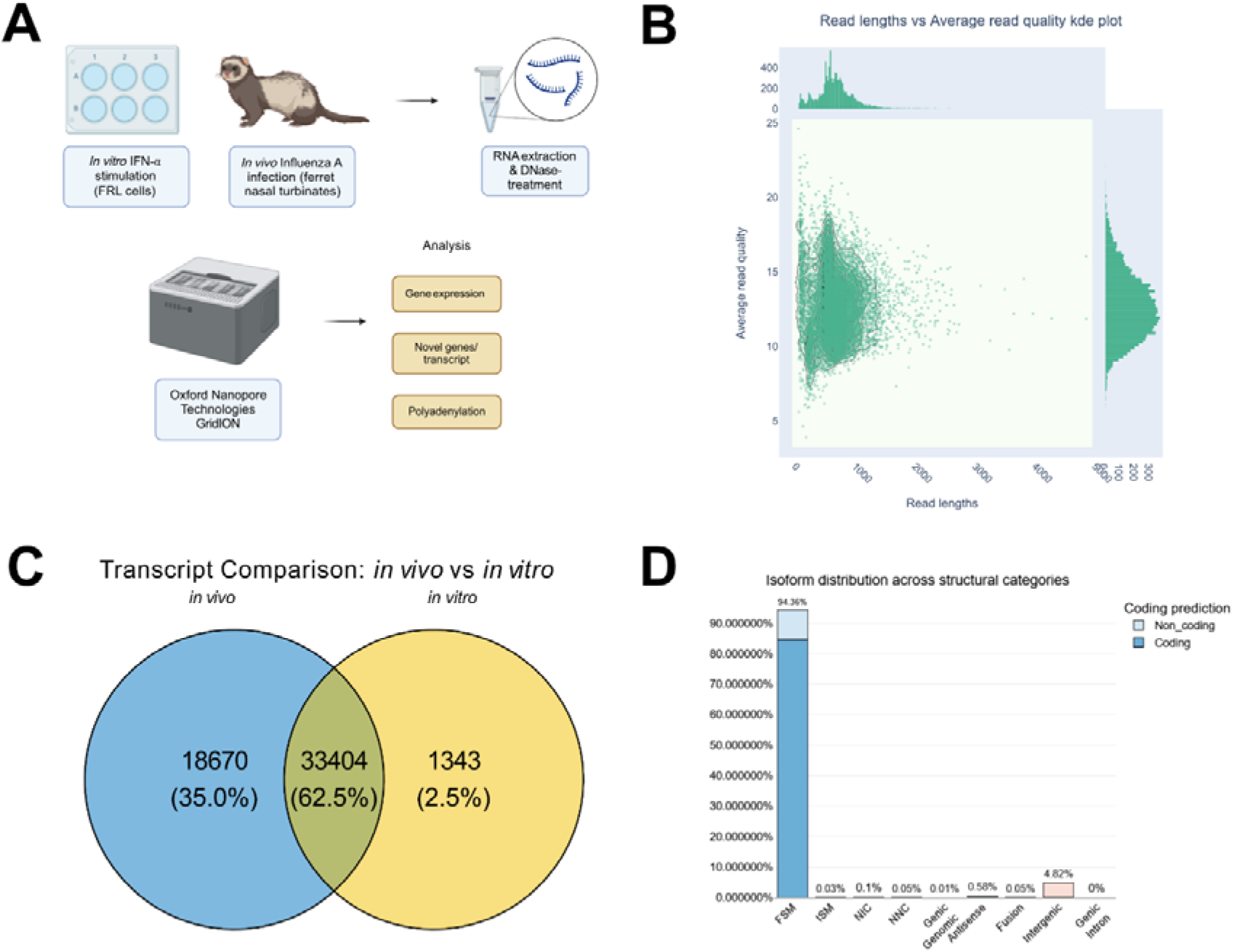
Characterization of ferret transcripts sequenced for this study. **A)** Flowchart of methods and analysis. Briefly, FRL cells were cultured in the presence or absence of recombinant ferret IFNα (*in vitro*) and domestic ferrets were mock-infected or infected via the intranasal route with IAV strain A/Perth/269/2009 (*in vivo*). Total RNA was extracted from the FRL cells 24 hour (hr) post treatment and nasal turbinate samples were collected from ferrets 6 or 24 hr post-infection (hpi), which were then treated with DNase. The RNA was sequenced on the ONT GridION using the SQK-PCB111.24 kit. Data analysis steps included gene expression, novel gene/transcript discovery and poly(A) analyses. **B)** Scatter kde plots for read lengths and average read quality derived from ONT PCR-cDNA sequencing. X-axis shows read-lengths in nucleotides and Y-axis shows average read quality. **C)** Venn Diagram showing the number of transcripts discovered in the *in vivo* vs *in vitro* datasets. **D)** *SQANTI3* isoform structures distributions across detected transcripts post-filtering via *SQANTI3*.

To visualize the ferret transcriptome through our long-read datasets we used a combination of *Bambu* and *SQANTI3* pipelines. After *Bambu* was used to detect novel transcripts and create a new annotation file, we checked the characteristics of the annotated reads using the *SQANTI3* ‘qc’ function (**s**). Of the final list of true isoforms, it was evident that more isoform models were able to be detected in the *in vivo* datasets compared with *in vitro* datasets (**Figure 1C**). Furthermore, most isoforms (∼94.36%) were determined to be of the Full-Splice Match (FSM) class, and some were of the intergenic class (∼4.82%) (**Figure 1D & Data S1,** see **Methods** for definition of transcript model classes). This indicates that most transcripts matched the splice junctions of existing reference transcripts and that some were found between genes, likely due to the previously unannotated novel genes. After final *Bambu* quantification, we observed novel (1,414), known (‘NM’: 64, ‘NR’: 0) and hypothetical (‘XM’: 57,475, ‘XR’: 10,942) transcripts, and 1,120 novel genes. These results highlight that many of the predicted transcripts can indeed be found *in vitro* and *in vivo* and the strength of utilizing ONT sequencing for novel transcript discovery.

### Identification and regulation of ferret genes orthologous to human ISGs

To determine if ferret orthologues of human ISGs with known anti-IAV activity had been upregulated (7), we performed differential expression analyses to compare mock/control samples to IFNα treated (*in vitro)* or IAV infected (*in vivo*) samples. Differential expression analyses revealed 1,120 significantly differentially expressed genes (DEGs) between mock- and IFNα-treated FRL cells (*in vitro*, padj < 0.05, **Figure 2A**). For *in vivo* datasets, we observed 20 and 1,131 DEGs at 6 (**Figure S1A**, padj < 0.05) and 24 hpi (**Figure 2B**, padj < 0.05), respectively, compared to mock controls. No ferret ISG orthologues of interest were upregulated at 6 hpi, suggesting that innate immune pathways require additional time for activation and expression of ISGs in IAV-infected ferrets. However, when examining *in vitro* and *in vivo* analyses at 24 hr post-IFN treatment or 24 hpi, respectively, we observed significant upregulation of many ferret ISG and IST orthologues of interest compared to the relevant mock controls (**Figures 2A-B, S1B-D**). The upregulation of a select group (n = 6) of ferret ISGs, which represent orthologues of human ISGs with known anti-IAV activity, was validated using RT-qPCR to examine induction in FLR cells cultured for 24 hr in the presence or absence of ferret IFNα. With the exception of *MOV10,* all ferret ISGs tested were significantly upregulated following IFN treatment (**Figure 2E**, p<0.01) and certain ISGs such as *Tetherin, ISG15, Viperin* and *TRIM22* were potently upregulated (300-500-fold compared to mock, **Figure 2D**). Ferret *OAS2* was moderately upregulated (∼8-fold compared to mock). Our previous study used colony PCR to confirm that the major transcript of ferret *Mx1* produced in FRL cells was XM_004762192.2 (8). Herein, Nanopore sequencing confirms this result (**Figure 2A**) and shows that this *Mx1* transcript is also strongly upregulated in the nasal turbinates of ferrets 24 hr after IAV infection (**Figure 2B**). Finally, we note that the trend in fold changes observed in RT-qPCR data (**Figure 2D**) correlates well with differential analysis data obtained following Nanopore sequencing (**Figure 2A**).

**Figure 2.**
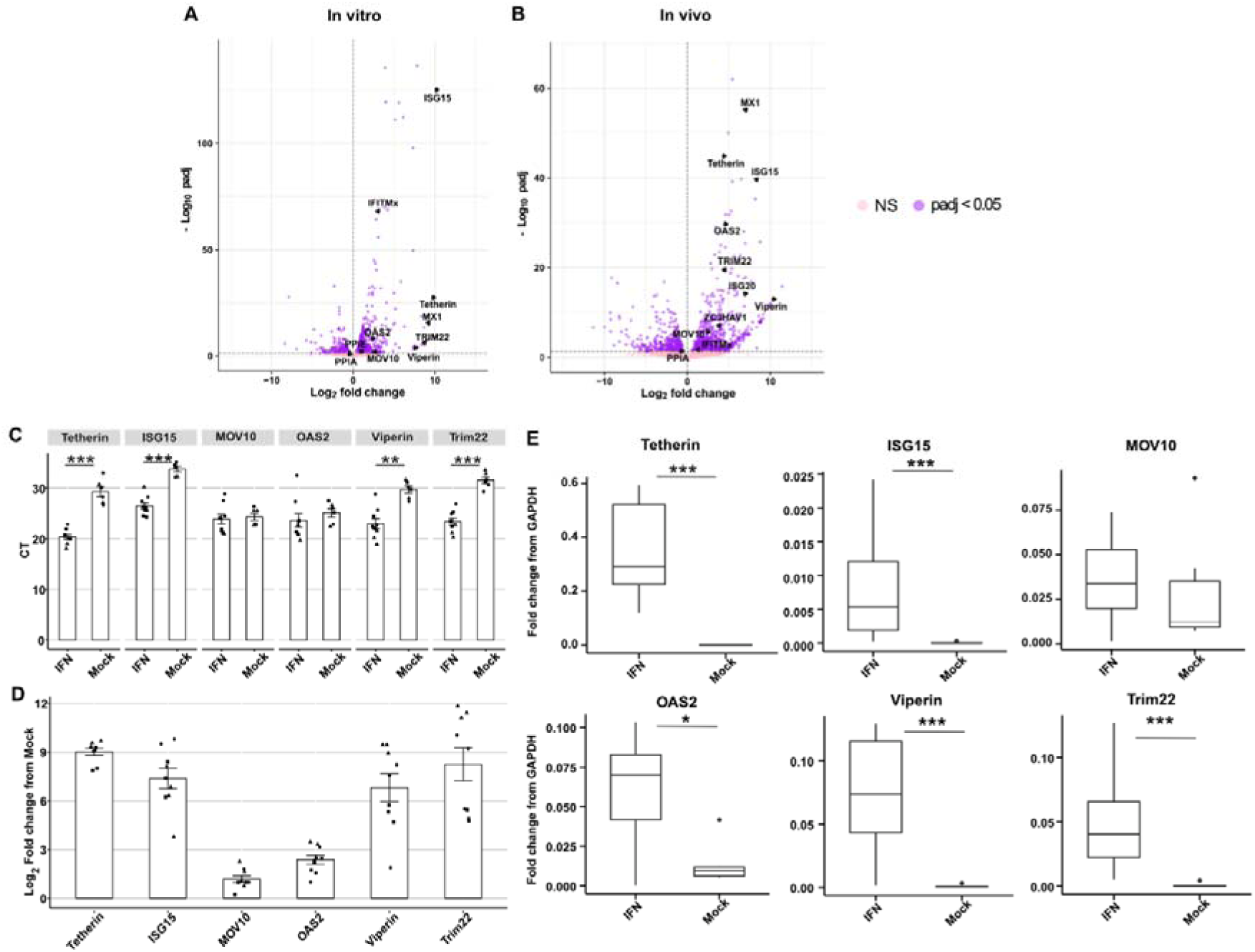
Upregulation of ferret genes orthologous to particular human ISGs in response to IFN treatment (*in vitro, 24 hr post-treatment*) or IAV infection (*in vivo, 24 hr post-infection*). Differential expression results in **(A)** *in vitro* and **(B)** *in vivo* datasets at the gene level. X-axis represents log_2_ fold changs and the Y-axis shows the −log_10_ p-adjusted value (padj), with a threshold of padj < 0.05, shown by the dotted horizontal line. **(C-E)** Six ISGs were validated following analysis of FRL cells cultured for 24 hr in the presence or absence of IFN. RT-qPCR results show **(C)** Raw CT differences between mock- and IFN-treated FRL cells**, (D)** Log_2_ fold change in expression levels relative to mock controls, calculated using 2^−ΔΔCT^ method, and **(E)** fold change relative to housekeeping gene *GAPDH* using the 2^−ΔCT^ method*. Pooled data from three independent experiments are shown, different symbols are used to denote each experiment. Statistical differences in CT and fold-change values from mock (C and E) were calculated using a mixed effects model. * p<0.05, ** p<0.01, ***, p<0.001*.

In addition to identifying ferret ISG transcripts that represent orthologues to human ISGs with known anti-IAV activity, we aimed to determine their similarity between different mammalian species. Therefore, we compared the predicted protein coding sequences of ferret ISGs identified in this study to published sequences of ISGs from humans, mice, and European mink, noting that mice represent the small animal model most commonly used to study IAV *in vivo* (**Figure 3**). While human and mouse genomes are well annotated, the European mink genome is incomplete and therefore predicted transcript sequences of mink ISGs were used in these analyses. Alignment results indicate that, with the exception of *OAS2*, all ferret ISGs have the highest degree of similarity to European mink. Dissimilarity between ferret and mink *OAS2* is likely artefactual and due to errors in annotation of the mink genome. *MOV10* and *Viperin* were the most conserved ISGs across different species (>80% similarity in protein sequence), while *Tetherin* showed the highest degree of divergence (**Figure 3**). In general terms, most ferret ISGs were to be closer to human than to mouse ISGs.

**Figure 3.**
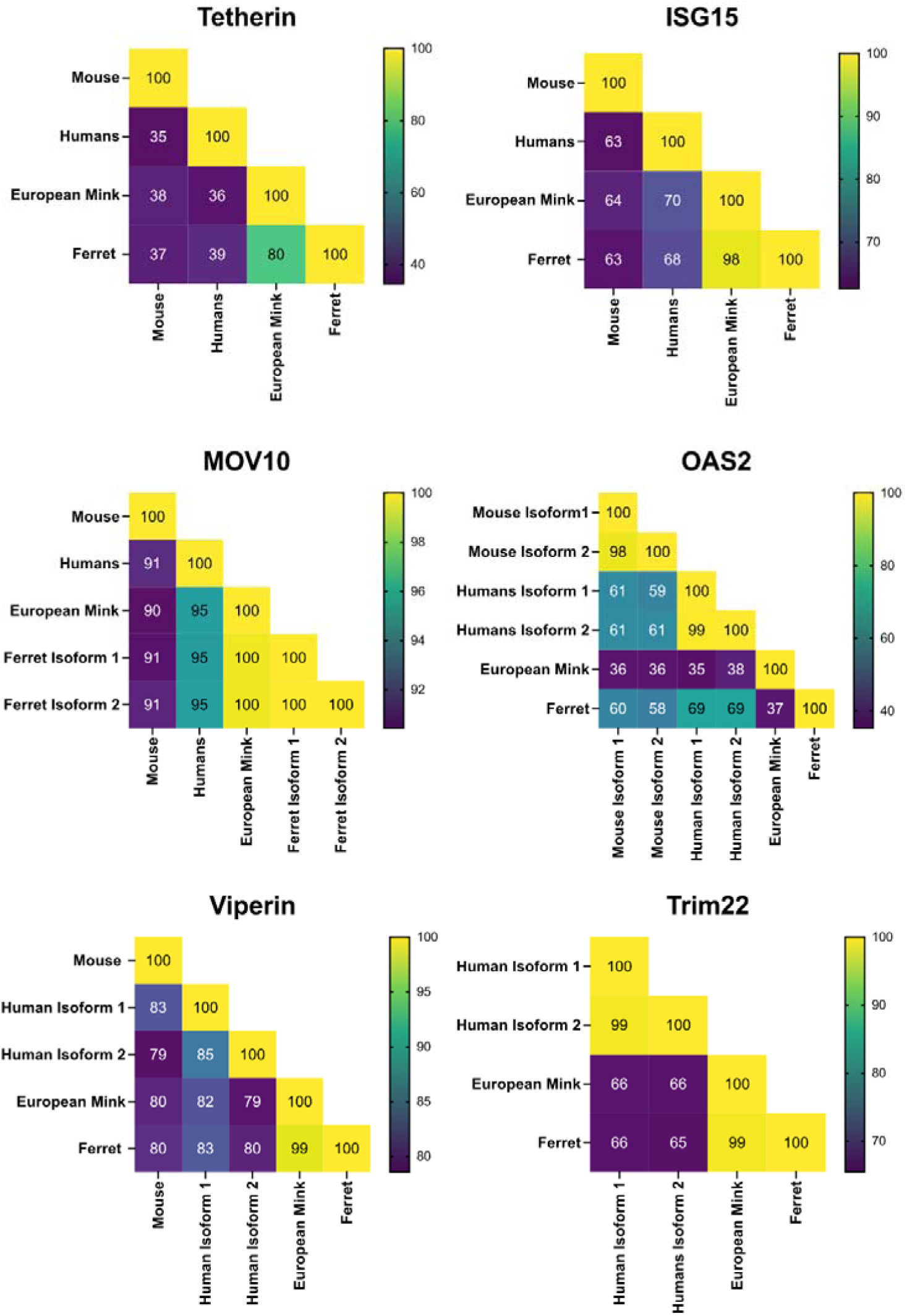
The protein coding sequence (CDS) for a select group of ISGs obtained from NCBI (for humans, mice and European mink (*Mustela lutreola*)) or from our data set (ferret), were aligned using *MUSCLE*. The percentage identity between each species is represented in a heatmap for each ISG. Of note, mice do not express an orthologue of TRIM22. For European mink, genome annotation remains incomplete and all protein CDS information was derived from predicted transcripts (denoted by XM accession numbers, see **Methods**). Moreover, we detected two isoforms of ferret MOV10, with isoform 1 representing the major species *in vitro* and isoform 2 *in vivo*. While minor isoforms of other ferret ISGs were detected, their expression levels were very low and they have therefore been excluded from this analysis.

The human ISGs *Tetherin, ISG15, MOV10, OAS2, Viperin* and *TRIM22* are very well characterized, and as such we know certain regions/sites/motifs are important for their protein localization, stability, signalling, antiviral activity, post translational modifications etc.(7). We therefore utilized our alignments (**Figure 3**) to interrogate whether these sites/regions/motifs were conserved between humans and ferrets, which can provide insights into whether these ISGs have similar function (and therefore similar antiviral activities) across the two species. Our results show that all major sites relevant to protein function and the antiviral activity of *ISG15* and *MOV10* are conserved between humans and ferrets, whereas differences were noted in some major sites between human and ferret *Tetherin, OAS2, Viperin* and *TRIM22*. (**Table 1**) These differences pertain to aspects of protein function such as NFκβ activation or enzyme activity (*Tetherin* and *OAS2*), protein localization (*TRIM22*) or protein-protein interactions (*Viperin*), highlighting species-specific differences in ISG structure which could be validated experimentally in future studies.

**Table 1.** Degree of conservation in regions/sites needed for protein function, signalling and/or antiviral activity (human amino acid numbering)

| Gene | Conserved Sites/Regions/motifs | Non-conserved Sites/Regions/motifs |
| --- | --- | --- |
| Tetherin | Sites needed for NFκβ activation (6Y, N65, N92, N123) and homodimer formation (C53, C63, C91) | Region needed for NFκβ activation (8-11) |
| ISG15 | Site needed for disulphide bond (C78) and signalling (Y96, R99, T101, T102, T103); LRLGG motif (152-187) |  |
| MOV10 | Region needed for interaction with AGO2 and APOBEC3g (995-998); DEAG box (678-681) |  |
| OAS2 | Sites needed for enzyme activity (D408+D410, Y421, D481, R544, K547) | Sites needed for enzyme activity (669-670) |
| Viperin | Sites needed for antiviral activity against Zika virus (K358), ubiquitination (K206), acetylation (K197) and protein stability (C83, C87, C90) | Leucine zipper motif (14-42) |
| TRIM22 | Sites needed for antiviral activity (C15, C18) | Nuclear localisation signal (257-275), Site important for formation of nuclear bodies (V943) |

### Characterization of novel ferret ISGs and ISTs

In addition to visualizing ferret orthologous of human ISGs, we also searched for novel unannotated ISGs and interferon stimulated transcripts (ISTs) in the datasets generated. We focussed our analyses on novel genes and transcripts (‘BambuGene’ and ‘BambuTx’) upregulated in the *in vitro* data set (IFNα stimulation, padj < 0.05), reasoning that these were most likely to represent potential ISGs and ISTs. Genes and transcripts of interest were (i) ranked via estimated average expression level in the IFN-treated datasets, and then (ii) selected for low, mid and high expression genes and transcripts depending on their rank position (**Figures 4A-B**). Through this process, we selected novel transcripts from three genes – *GCA*, *LIPA* and *LOC106005667*. The novel ISTs showed exon-skipping events and alternative 5’ ends compared to the annotated transcripts (**Figure 42B**). The selected ISGs and ISTs were then validated using PCR and RT-qPCR to confirm a trend towards higher expression in datasets from IFN-treated compared to mock-treated FRL cells for most ISGs/ISTs, although this was not significant (**Figures 4C-E**). Additionally, the novel gene *BambuGene8284* appeared to show remarkably high expression levels in datasets from both mock and IFN-treated FRL cells (**Figures 4F**), which further validated the existence of the transcripts. Analysis of gene expression data from sequencing showed that this gene also was upregulated in the nasal turbinates of IAV-infected ferrets, and expression levels increased markedly between 6 hpi and 24 hpi (**Figure S2**). Several other *BambuGenes* were also upregulated following IFN treatment (*in vitro*) or IAV infection (*in vivo*) (**Figure S2-3**), and future studies will determine their coding potential and their role in modulating IAV infection. Overall, our analysis reveals the presence of novel ISGs and ISTs that have not yet been annotated in the reference ferret genome and transcriptome.

**Figure 4.**
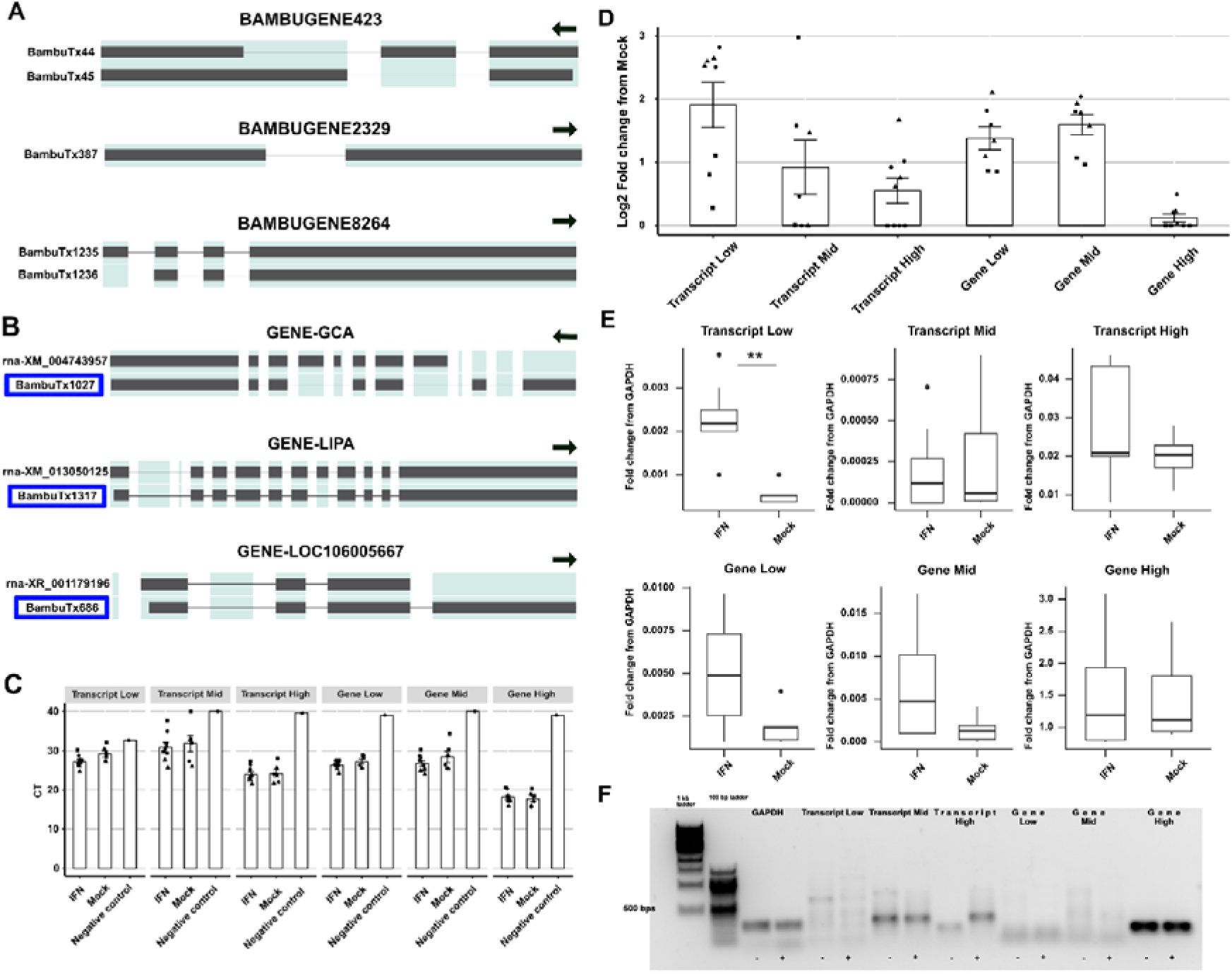
Novel ISGs and ISTs derived from *Bambu* analysis. **A)** Novel ISGs in order of expression - *BambuGene423* (low) *BambuGene2329* (mid), and *BambuGene8264* (high). *BambuGene8264* revealed exceptionally high expression compared with the other novel ISGs with two main novel isoforms. Arrows indicate the direction of the strand from 5’ to 3’. **B)** Novel ISTs in order of expression - GCA – *BambuTx1027* (low), *LIPA* – *BambuTx1317* (mid), and *LOC106005667* – *BambuTx686* (high). Novel transcripts from known genes are highlighted in blue boxes. Only reference transcripts with the highest expression are shown for comparison. Arrows indicate the direction of the strand from 5’ to 3’. **C)** Raw CT values of novel ISTs and ISGs from RNA extracted from mock- or IFN-treated FRL cells. Negative controls (primer only) have been included for comparison. **D)** Log fold change in expression levels relative to mock controls, calculated using 2^−ΔΔCT^ method, and **E)** fold change relative to housekeeping gene *GAPDH* using the 2^−ΔCT^ method. Pooled data from three independent experiments are shown, different symbols are used to denote each experiment. Statistical differences in CT and Fold change value from mock (C and E) were calculated using a mixed effects model. * p<0.05, ** p<0.01, ***, p<0.001. **F)** Representativ**e** DNA gel to visualise qPCR products from *n vitro* samples.

### Novel trans-spliced paralogous reads in potential ferret *IFITM* genes

The *IFITM* (Interferon-induced transmembrane protein) gene family encodes for proteins which mediate antiviral activity, including against IAV, by limiting the early stages of the virus replication cycle (27). In humans, three IFITMs (IFITM1, 2 and 3) have been particularly well characterised and are known to localize to plasma (IFITIM1) or endosomal membranes (IFITM2 and 3) where they can inhibit entry by viruses that enter cells by direct fusion (IFITM1) or by endocytosis (IFITM2 and 3) (28). In contrast, ferret IFITM genes are generally not well characterized and the transcript sequences available are predicted from short-read data in NCBI. A recent study used qPCR to demonstrate that the hypothetical (i.e.‘XM’) ferret IFITM transcripts were upregulated in IFN-treated and IAV-infected FRL cells (29). Herein, we also observed similar results in both *in vitro* and *in vivo* datasets (**Figures 2A**-**B**). Compared to the well-characterized human transcriptome (**Figure 5A**), the annotated hypothetical transcripts in ferrets revealed greater variety in potential orthologous regions (**Figure 5B**). Furthermore, upon visualization using the *Integrative Genomics Viewer (IGV)*, we noted reads which mapped to exons of different hypothetical (i.e. ‘XM’) transcripts of *IFITM* genes, suggesting trans-splicing events (**Figure 5B**). We also observed evidence of reads which showed additional 5’ exons compared with the annotated exons of genes (**Figure 5C**), implying an incomplete annotation of the *IFITM* transcripts. To understand the functional consequences of trans-spliced transcripts and transcripts with extra 5’ exons (denoted Transcript 1, Transcript 2 and Transcript 3 in this manuscript), we predicted their protein structures. The resulting predicted proteins all contained a conserved CD225 sequence that regulates vesicular membrane fusion, contributing to the most important functional domain for IFITM, as also observed in the reference *IFITM* transcripts (**Figure 5D**) (30). The major sequential differences between the three predicted proteins appeared to be in the N-terminal region.

**Figure 5.**
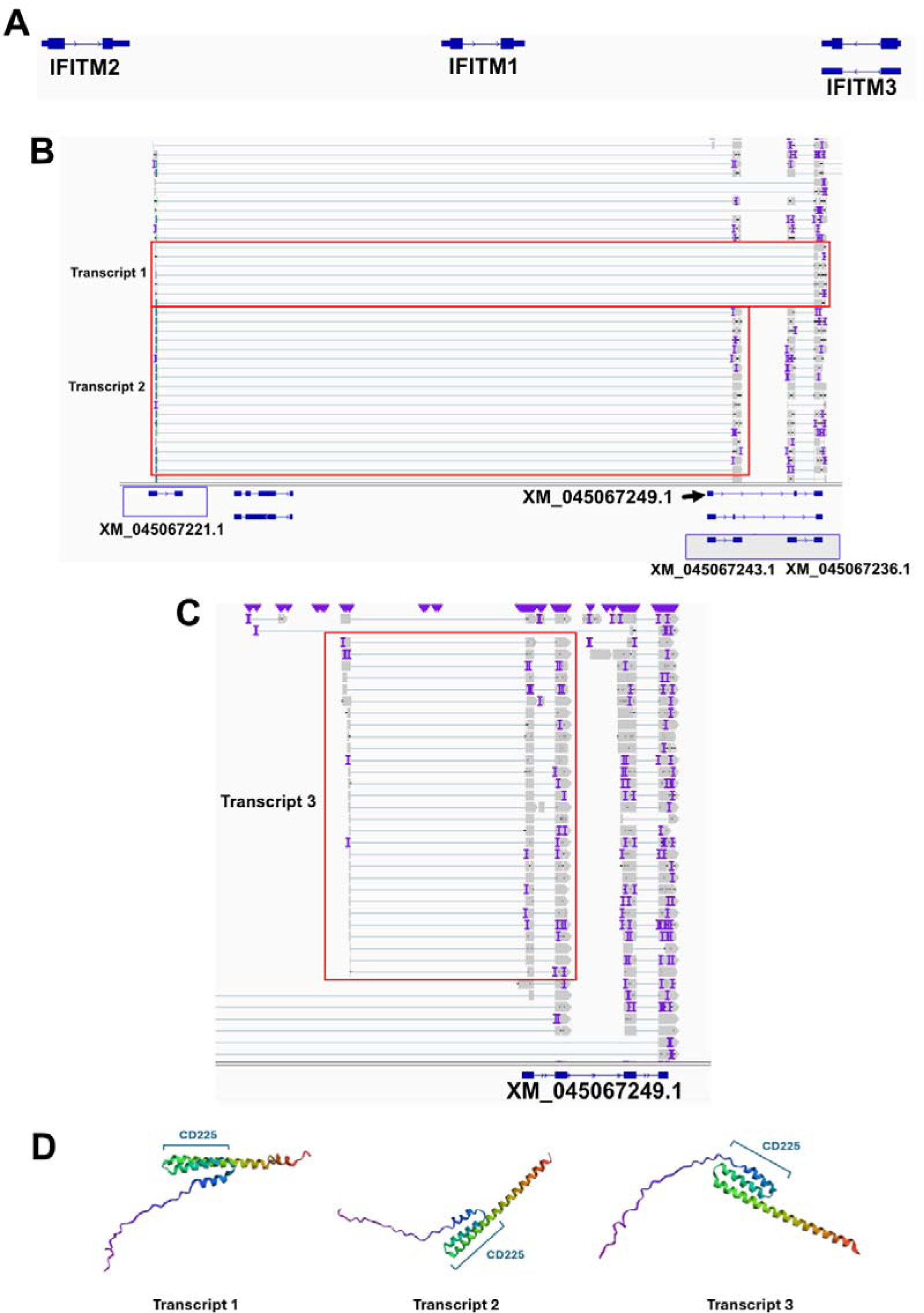
Potential novel trans-spliced transcripts in the ferret *IFI TM* family of genes. **A)** *IGV* plot visualization of human *IFIT M1-3* reference transcripts. **B)** *IGV* plot visualization of putatively trans-spliced *IFITM* transcripts (transcripts 1 & 2). The relevant reads are highlighted via a red box. Reference transcripts that appeared to be involved in the trans-splicing events are highlighted via the blue boxes. **C)** *IGV* plot visualization of reads suggestive of extra 5’ exons in putative ferret *IFITM* transcripts (transcript 3). **D)** Protein structure estimates of trans-spliced transcripts using *trRosetta*.

To investigate whether trans-spliced *IFITM* transcripts were actually expressed *in vitro* and/or *in vivo,* we first designed primers specific to each novel trans-spliced transcript (taking care to ensure the primers were specific; **Figure S4**). We then utilized RT-qPCR and RNA extracted from mock- or IFN-treated FRL cells, or mock- or IAV-infected nasal turbinates of ferrets to determine expression levels. (**Figure 6**). In our *in vitro* dataset, we noted that a transcript sequence in NCBI labelled as ferret *IFITM3* (XM_045067236.1) was highly upregulated in FRL cells following IFN treatment. However, as this transcript exhibits a high degree of sequence similarity to both human *IFITM2* and *IFITM3*, we subsequently refer to this as *IFITMx* in our studies. Due to its high expression levels, we utilised *IFITMx* as a positive control in the RT-qPCR assays below. Compared to negative controls, RT-qPCR results from FRL cells demonstrated an appreciable difference in CT values for *IFITM* transcript 1, but not transcripts 2 and 3, suggesting they are not being expressed at levels detectable by RT-qPCR (**Figure 6A**). In contrast, *IFITMx* was readily detected by RT-qPCR. Interestingly, none of the ferret *IFITMs* were expressed at high levels relative to the housekeeping gene *GAPDH* (**Figure 6C**), even following IFN stimulation of FRL cells, although it should be noted that *IFITMx* expression did increase (∼5-fold relative to mock) in response to IFN treatment. Visualization of the RT-qPCR products showed a distinct band for *GAPDH* and a faint, but distinct, band for *IFITMx* which increased in intensity following IFN treatment. However, no clear bands could be observed for any of the novel *IFITM* transcripts (**Figure 6E**). The trends in CT and fold change derived from *in vivo* samples were similar to those from i*n vitro* samples, with *IFITMx* showing a ∼23-fold increase relative to mock in IAV-infected samples, noting that this was not significant (**Figure 6B & D**). Visualization of RT-qPCR products showed a distinct band for *GAPDH* and *IFITM* transcript 1, but not for *IFITM* transcript 2 or 3 (**Figure 6F**). Together, these results suggest that *IFITM* transcript 1 is produced, with stronger evidence of this trans-spliced transcript is produced *in vivo* although it does not appear to be upregulated by IFN treatment (*in vitro*) or IAV infection (*in vivo*).

**Figure 6.**
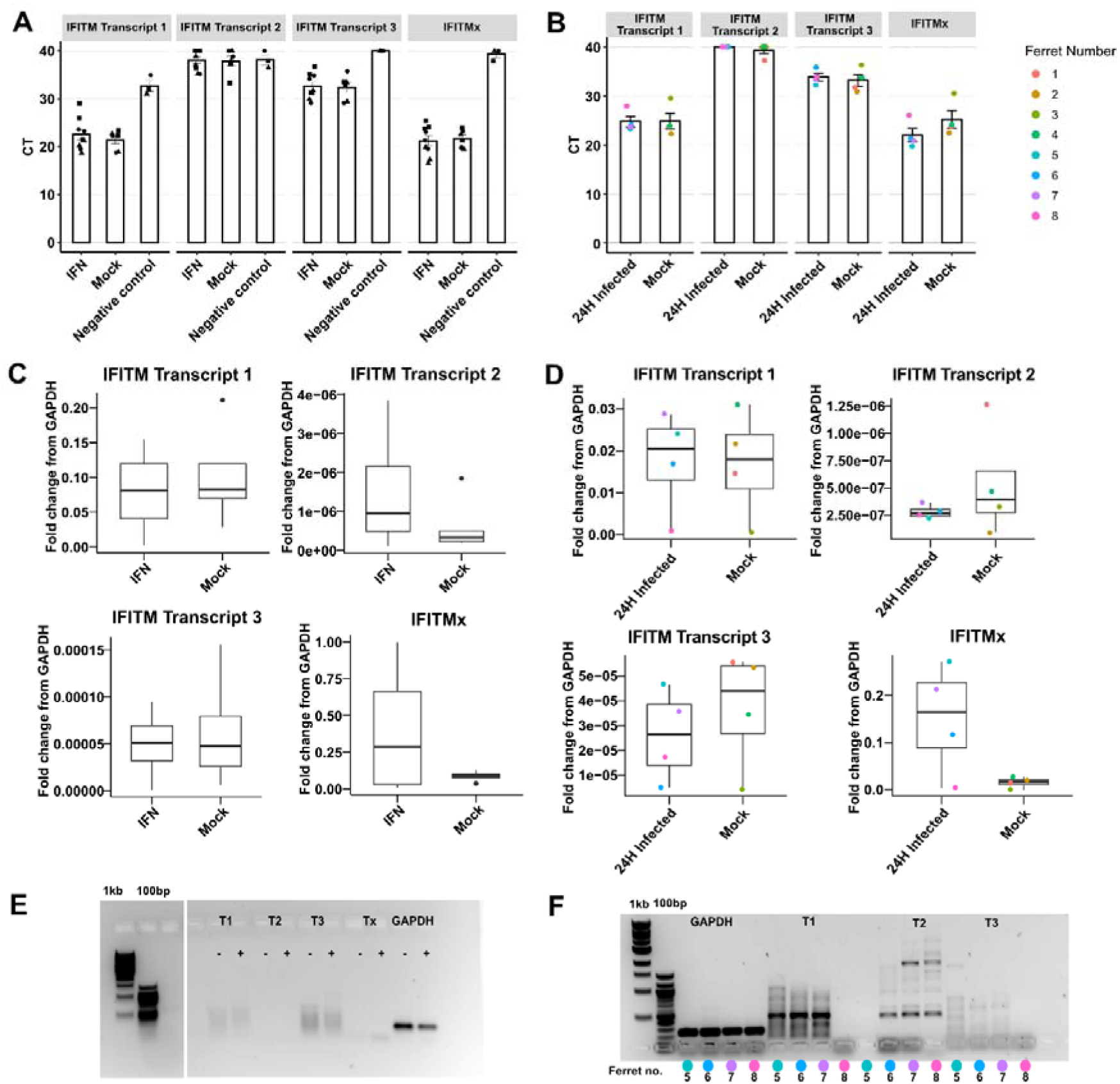
Validating the expression of novel trans-spliced *IFITM* transcripts by RT-qPCR. (**A-B)** Raw CT values of *IFITM* transcripts under different experimental conditions in FRL cells **A)** or in ferret nasal turbinates **B)**. **(C-D)** Expression level of *IFITM* transcripts under different experimental conditions in FRL cells **C)** or ferret nasal turbinates **D)**, with fold change relative to housekeeping gene *GAPDH* calculated using the 2^−ΔCT^ method. For *in vitro* data, pooled data from three independent experiments are shown (**A/C**), and different symbols are used to denote each experiment. For *in vivo* data, different colors indicate individual animals. Statistical differences in CT and fold change values from mock was calculated using a mixed effects model. * p<0.05, ** p<0.01, ***, p<0.001. **(E-F)** Representative DNA gels to visualise qPCR products from *in vitro* **E)** or *in vivo* **F)** samples.

### Ferret poly(A) tail regulation is associated with viral infection pathways

A favorable feature of ONT RNA-seq is the ability to estimate transcriptome-wide poly(A) tail length. As such, we investigated the lengths of the poly(A) tails of ferret mRNA. All datasets, despite having varied conditions, showed comparable distributions (**Figure 7A**). The ferret transcriptome also had a median of ∼52 nt in the mitochondrial genes and ∼68 nt in the nuclear genes, which agrees with studies in human polyadenylomes (14,16,17) (**Figure 7B**). The *in vivo* dataset (IAV-infected ferrets at 24 hpi) also exhibited the highest proportion of reads with poly(A) tails greater than 400 nt (**Table 2**), which suggests a correlation between infection and poly(A) tail elongation, in line with previous work in human cell lines (14). To determine if the poly(A) tail length varied depending on Kyoto Encyclopedia Genes and Genomes (KEGG) pathways, we used the Gene Set Enrichment analysis tool to identify pathways enriched for shorter or longer tails in mock/control conditions only. Interestingly, two main groupings were apparent based on the maximum peak in density of poly(A) tails; a smaller subset of pathways with shorter maximum densities below ∼70 nt, and a larger group of pathways with longer maximum densities (**Figures 7C-D**). This indicates that as in other species, the ferret polyadenylome appeared to have a distinct set of genes which harbor shorter or longer poly(A) tails depending on their functions. Notably, both mock-treated FRL cells (**Figure 7C**) and mock-infected ferret nasal turbinates (**Figure 7D**) showed the *proteasome* pathway as being part of the shorter length poly(A) group and other metabolic pathways. Additionally, differential polyadenylation analysis revealed that IFN-treated FRL cells had elongation of poly(A) tails in the mRNA from genes involved in the *ribosome*, *coronavirus disease 19* and *oxidative phosphorylation* KEGG pathways, in comparison to mock-treated FRL cells, revealing a potential relationship between poly(A) tail elongation and innate immune responses in domestic ferrets (**Figures 7E-F**). Overall, the polyadenylome of ferrets align with the human polyadenylome (16), showing its feasibility as a model organism and as a proxy for understanding human responses to IFN treatment or IAV infection.

**Figure 7.**
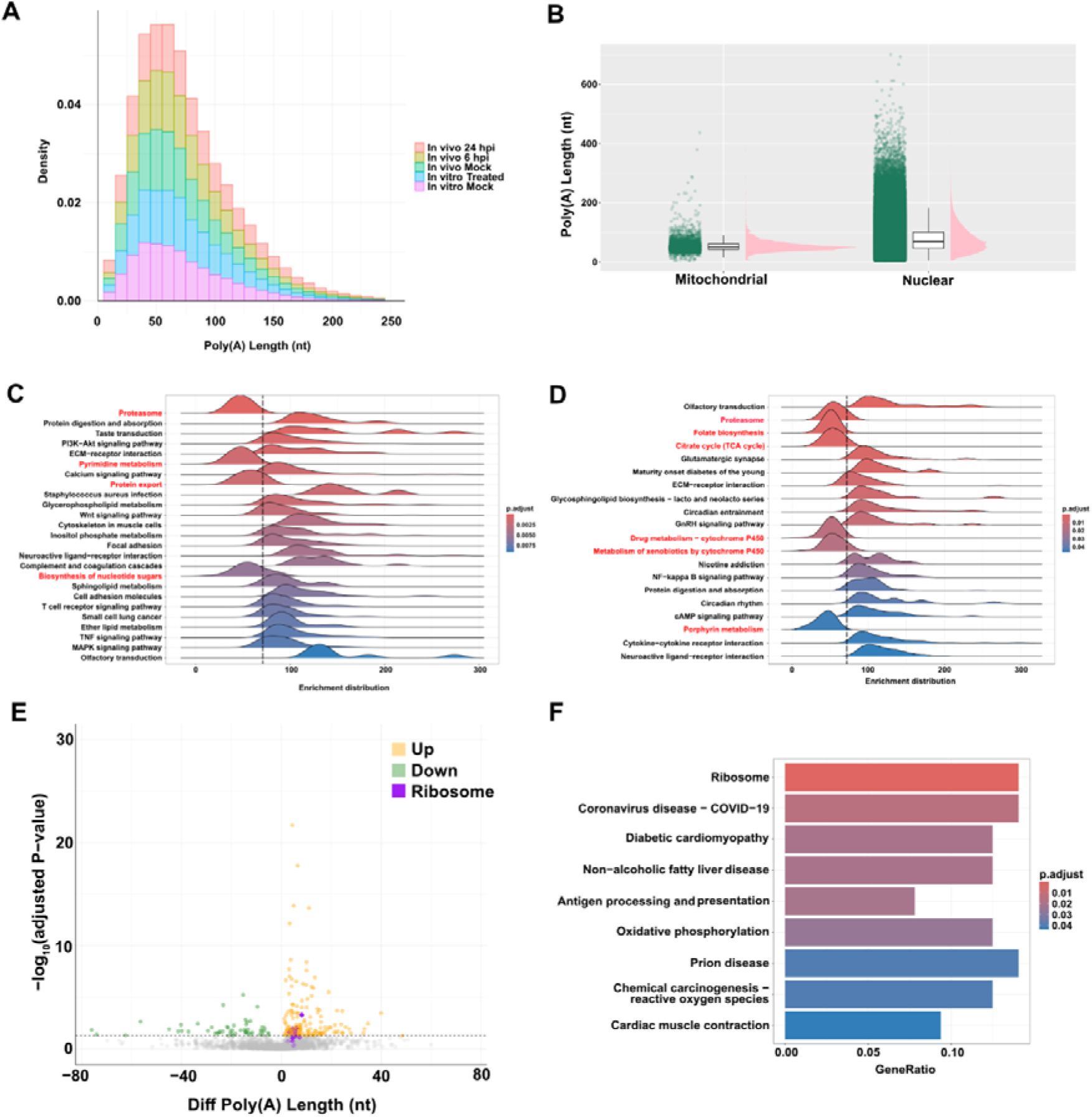
Polyadenylation states in the ferret transcriptome. **A)** Density distributions of poly(A) lengths in each dataset. Only nuclear RNA has been included. **B)** Poly(A) distributions between nuclear and mitochondrial RNA. A random sample of 300,000 reads across the different datasets were used. **(C-D)** KEGG pathway GSEA results using **C)** *in vitro* mock-control and **D)** *in vivo* mock-control data. Pathways highlighted in red are those in which the maximum density peaks are deemed with lower average poly(A) length (indicated by the dashed vertical line). The genes were ranked based on poly(A) lengths. **E)** Differential polyadenylation results between *in vitro* 24 hpi IFN-treated vs mock-control data. X-axis shows the difference in maxpeak poly(A) length between the two datasets compared and the Y-axis shows the −log_10_ padj, with a threshold of padj < 0.05. Each dot represents a gene, where green = decreased polyadenylation, yellow = increased polyadenylation and purple = genes involved in the *ribosome* KEGG pathway, shown to be increased in polyadenylation. **F)** KEGG pathway enrichment analysis of genes with increased poly(A) tail lengths upon IFN-treatment compared with mock-control in the *in vitro* dataset with a threshold of padj < 0.05.

**Table 2.** Number of reads belonging to each dataset and reads with long poly(A) tails (>400 nt).

| Dataset | Total Read Count | #Reads with Poly(A) > 400 nt | Proportion of Reads with Poly(A) > 400 nt |
| --- | --- | --- | --- |
| In vitro-mock | 5,309,690 | 729 | 1.37E-04 |
| In vitro-IFN treated | 5,026,980 | 562 | 1.12E-04 |
| In vivo-mock | 4,982,784 | 1,095 | 2.20E-04 |
| In vivo-IAV infection, 24 hpi | 11,395,103 | 4,270 | 3.74E-04 |
| In vivo-IAV infection, 6 hpi | 12,330,668 | 3,120 | 2.53E-04 |

## Discussion

The European domestic ferret (*Mustela putorius furo*) reference genome and transcriptome have been curated predominantly through computational predictions, despite the ferret being a benchmark species for IAV-infections (31). This was clearly apparent due to the proportion of known versus hypothetical transcripts detected via our *Bambu* analysis which identified known (‘NM’: 64, ‘NR’: 0) and hypothetical (‘XM’: 57,475, ‘XR’: 10,942) transcripts. This makes investigating ferret responses to different stimuli using multi-omic approaches very challenging. In this study, we comprehensively evaluated the ferret transcriptome using ONT long-read RNA-sequencing, which validated the annotated ferret transcripts in NCBI, as well as detecting novel transcripts through our reference-guided analyses. We detected over 1,400 novel transcripts, confirming the importance of utilizing long-read technology to assess an incomplete transcriptome. We also observed increased diversity in transcripts detected in the *in vivo* data (**Figures 1C, S1B&D**), which may be caused by the greater variation between individual ferrets compared with the homogenous FRL cell line. Moreover, the nasal epithelia is composed of heterogenous cells, including ciliated, secretory and basal cells (32), as well as a varied composition of immune cells both in homeostasis and during infection, contributing to the increased diversity. We note that assessing the bulk response of these cell populations by averaging across cell-types will mask the intricacies of cell-type-dependent transcriptional regulation. Therefore, further research should be conducted to assess the IST expression at a single-cell level in the ferret nasal epithelia.

Previous studies investigating ferret ISGs in the context of viral infections have been hampered by two main challenges: (i) the inability to determine which ferret ISGs are genuinely expressed during viral infection, and (ii) the existence of multiple predicted transcript variants for each ISG in the NCBI database, with no indication of which isoform represents the major transcript produced during infection or in response to IFN stimulation (8,29). For example, in our previous study on ferret Mx proteins we were unable to characterize ferret *Mx2* at the protein level due to the presence of 14 predicted transcript variants on NCBI, making it difficult to identify the biologically relevant isoform for functional studies. Similarly, a prior study aiming to characterize ferret *IFITMs* first required annotation of the transcript sequences from the gene locus before RT-qPCR validation of their expression could be performed (29). With these limitations in mind, the first aim of this study was to identify the major transcript variants of ferret ISGs that are orthologous to human ISGs with known anti-IAV activity. We successfully identified the dominant transcript variants of ferret *Mx1* (previously characterized in (8)), *Tetherin, ISG15, MOV10, TRIM22, Viperin,* and *OAS2*. Using RT-qPCR, we demonstrated that all of these ferret ISGs, except *MOV10*, were upregulated following IFN treatment. Unfortunately, we were not able to determine the major transcript variant of ferret *Mx2*, due to this transcript being very long, and poor sequencing coverage in the N-terminal region where most variation between the transcript variants are observed.

Of note, the initial ferret *OAS* transcript identified (XM_004753413.2, annotated as *OAS3* on NCBI) in this analysis appeared unusually long and resembled a fusion between human *OAS2* and *OAS3*. Upon further investigation, we determined that there was stronger evidence of a shorter transcript being present, corresponding to region 3456-6750 of XM_004753413.2. This shorter transcript showed a high sequence similarity to human *OAS2*. Accordingly, subsequent RT-qPCR analysis and protein alignment studies focused on this transcript, which we refer to as ferret *OAS2* in this manuscript. Notably, there is currently no ferret *OAS2* annotated in the NCBI database, making this a novel finding. Similarly, we identified a ferret *IFITM* sequence (XM_045067236.1) which has not been described in the previously published Horman *et al* paper (29) and has a sequence similar to both human *IFITM2* and *IFITM3*. We have denoted this as *IFITMx* in this manuscript. In addition, we have some evidence to indicate that trans-spliced *IFITM* transcripts are produced in ferrets, which were most prevalent in IAV-infected ferrets and were not strongly induced by IFN treatment in FRL cells. While protein structural analysis suggests that these trans-spliced transcripts can encode for functional proteins, whether they play a role in virus infections remains to be determined. Trans-splicing events can lead to novel protein formation, creating neoantigens which may elicit a varied or greater immune response. Otherwise, the trans-spliced transcripts can act as regulatory non-coding RNA and participate in increasing the genome complexity. Taken together, while these transcripts were rather poorly expressed in comparison with reference *IFITM* transcripts, we hypothesize that the trans-spliced *IFITM* transcripts may act as back up transcripts to encumber diversity within *IFITMs*, to act against IAV infections.

In addition to identifying the dominant transcript variants of ferret ISGs, we were able to compare the protein coding region of the ferret ISGs with other mammalian species, such as humans, mice and European mink. As expected, there was a high degree of similarity between ferret and mink ISGs, noting that the mink database is also incomplete. This incompleteness likely played a role in the high degree of divergence observed between mink and ferret *OAS2*. Regardless, the information gleaned from this study and future studies characterizing the antiviral activity of ferret ISGs in more detail, are likely to provide additional insights into mink ISGs as well.

In addition to comparing protein coding sequences, our alignment data also provided us with some potential insights into differences in protein function and/or localization between ferret and human ISGs. For example, data from **Table 1** of this study indicate that ferret *TRIM22* contains mutations within the nuclear localization signal present in human *TRIM22*. As a result, unlike its human counterpart, ferret *TRIM22* may not localise to the nucleus during infection (33). Further experiments are needed to confirm this hypothesis and determine if ferret *TRIM22* has similar antiviral activity to human *TRIM22* or not.

In addition to identifying orthologues of known ISGs, we were able to find unannotated ISGs and ISTs which were upregulated following IFN treatment of FRL cells and/or IAV infection of ferrets (**Figures S2-3**), with *BambuGene8284* showing impressively high baseline expression levels *in vitro* and *in vivo*. While this novel gene does not align to many of the known ISGs in humans, *Blastn* analysis suggests that this gene encodes for a noncoding RNA, which is of interest as ncRNAs are increasingly recognized as ISGs with the potential to modulate viral infection (34). Further work will be required to fully elucidate the characteristics of these novel ISGs and ISTs.

Polyadenylation is an important co-transcriptional modification which occurs not only in mammals but also in viruses (10,35) and plants (36). According to our previous studies on the polyadenylome of human transcriptome (14,16), we found a correlation between the ferret polyadenylome and the human polyadenylome in terms of general distribution and effect during an event such as IAV infection and/or IFN-treatment. Particularly, IFN-treated FRL cells exhibited increased poly(A) tail lengths of mRNA of genes related to KEGG pathways commonly associated with viral or bacterial infection, such as *ribosome*, *COVID-19* and *oxidative phosphorylation* (**Figures 7E-F**). This is in alignment with our previous results with Calu-3 cells infected with SARS-CoV-2 (14). Poly(A) tail length has been correlated with the stability of the mRNA transcript (37), and suggests that this mechanism is part of the anti-viral arsenal of the host or contributes to the viral manipulation of host ribosome functions. However, no significant pathway enrichment was found in the *in vivo* datasets. This suggests i) a host-specific activity of the polyadenylome, ii) the species-dependent response to viral infection, or iii) differences in responses to viral infection vs IFN-treatment. Extending this work to determine the transcriptomic changes in IAV infection vs IFN-treatment in both *in vitro* and *in vivo* settings would be beneficial to understand these differences, in addition to understanding the aforementioned expression-level differences.

To date, only two studies have examined the ferret transcriptome following viral infection. One study sequenced and annotated the ferret transcriptome in lung lobes and lymph nodes after IAV infection (38), while another investigated transcriptomic responses in ferret lungs following henipavirus infection (39). The earlier study by León *et al.* (2013) was particularly instrumental in developing the ferret genome, with many current NCBI transcript annotations derived from that work. However, both studies used Illumina short-read sequencing technology, which limits the ability to resolve full-length transcripts and detect transcript isoforms. Furthermore, both focused exclusively on lung tissue, despite the nasal epithelium being a key site of IAV replication, and a site whose transcriptomic response remains largely unexplored.

To address these gaps, we conducted long-read sequencing on ferret lung cells ± IFN treatment (*in vitro*), as well as ferret nasal turbinates ± IAV infection (*in vivo*). While a recent study (40) used long-read sequencing to characterise the ferret immune repertoire, focusing on IgG and T cell receptor transcripts in splenocytes and lymph nodes, our study is, to our knowledge, the first to apply this technology to profile early innate immune responses in ferret nasal tissues. A limitation of our study was the lower RNA integrity (RINe) scores in the nasal turbinates samples, due to the sample collection method, which likely contributed to shorter N50 values. Nonetheless, we achieved sufficient coverage to capture full-length transcripts. Furthermore, due to the PCR amplification of our transcripts, only relative quantification was able to be carried out due to potential PCR bias. While Unique Molecular Identifiers (UMIs) were incorporated into the protocol to collapse the PCR duplicates, the UMI detection rate was insufficient to perform the deduplication, which may be caused by the higher error rates of the early version of the PCR cDNA barcoding kit. Despite this, amplified data is beneficial for detecting novel transcripts, which was the key mission of this study. This work may be extended by utilizing the most recent version of the ONT kit or direct RNA-sequencing, leading to higher accuracy and an enhanced view of the ferret transcriptome. Overall, our study represents the first long-read transcriptomic profiling of early antiviral responses in ferret respiratory tissues and provides an essential resource for improving gene annotation, understanding ISG expression dynamics, and supporting future functional studies in this important animal model.

## Methods

### Cells and viruses

Ferret lung epithelial (FRL) cells, an adenovirus 5-immortalized cell line (kindly provided by Prof. Tuck-Weng Kok, University of Adelaide), were maintained and passaged in Dulbecco’s Modified Eagle Medium (DMEM) (Gibco) supplemented with 10% (v/v) FBS, 2 mM L-glutamine (Gibco) and 1 mM sodium pyruvate (Gibco). Madin-Darby Canine kidney (MDCK) cells (ATCC CCL-34) were maintained and passaged in RPMI 1640 medium supplemented with 10% (v/v) fetal bovine serum (FBS, Gibco), 2 mM L-glutamine and 1 mM sodium pyruvate.

The seasonal IAV strain used in this study A/Perth/265/2009 (Perth/09, H1N1pdm09) was obtained from the WHO Collaborating Centre for Reference and Research on Influenza (WHO CCRRI), Melbourne, Australia. Viruses were propagated in the allantoic cavity of 11-day embryonated chicken eggs following standard procedures [31] and titres of infectious virus were determined on MDCK cells by standard plaque assay and expressed as plaque-forming units (PFU) per mL [32].

### Detection of ferret ISGs by qPCR following in vitro induction

To test for induction of ferret ISGs *in vitro*, FRL cells were seeded in 6-well plates 24 hr prior to treatment with 100 ng/µL of recombinant ferret IFNα (a kind gift from Tim Adams, CSIRO Manufacturing, Parkville, Australia). At 24 hr post-treatment, total RNA was extracted using the RNeasy Plus Mini Kit (Qiagen) and treated with amplification-grade Dnase I (Sigma Aldrich) and RNA concentration was then standardized across samples. RNA was then converted to cDNA using the SensiFAST cDNA Synthesis Kit (Bioline). SYBR green-based qPCR was used to determine the expression of ferret genes relative to housekeeping gene *GAPDH* (glyceraldehyde 3-phosphate dehydrogenase) using the SensiFAST SYBR kit (Bioline). Data acquisition was performed using the QuantStudio 7 Flex Real-Time PCR System (Applied Biosystems). For visualisation, qPCR products were run on an 0.8% agarose gel. Primer sequences used for qPCR are as below:

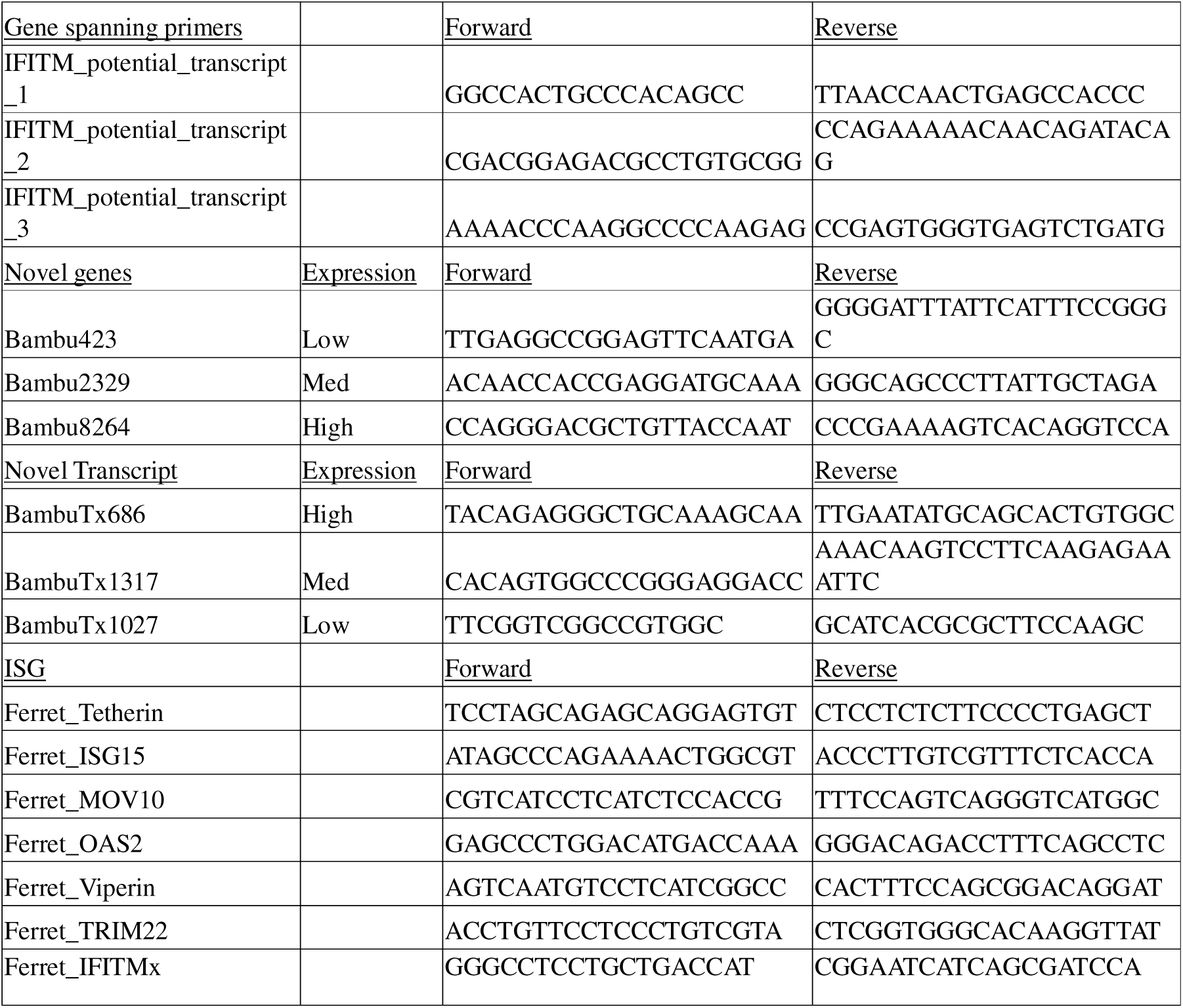

### Ethics statement

Ferret experiments were conducted with approval from the University of Melbourne Biochemistry & Molecular Biology, Dental Science, Medicine, Microbiology & Immunology, and Surgery Animal Ethics Committee (AEC# 20033), in accordance with the NHMRC Australian code of practice for the care and use of animals for scientific purposes.

### IAV infection of ferrets

Adult outbred ferrets (600-1500 gram) were housed in the Bioresources Facility at the Peter Doherty Institute for Infection and Immunity, Melbourne, Australia. Prior to the commencement of experiments, hemagglutination inhibition assays were used to confirm all animals to be seronegative against IAV strain A/Perth/265/2009 (Perth/09, H1N1)pdm09). For IAV infection, 12 ferrets were anesthetized (25 mg/kg ketamine and 5 mg/kg ilium Xylazilin a 1:1 (vol/vol) mixture) and 8 were inoculated by dropwise intranasal delivery of 500uL of PBS containing 10^7^ PFU of Perth/09 and 4 were mock-infected with an equivalent volume of PBS. At either 6 or 24 hr post-infection, ferrets were euthanized for collection of nasal tissues. For euthanasia, ferrets were anesthetized using a mixture of ketamine and xylazine following pentobarbitone sodium (Lethabarb Troy Laboratories) injection. Sections of nasal tissues were stored in 5 mL RNALater overnight at 4°C and tissues were then frozen at −80°C. Total RNA was extracted from ferret nasal tissue samples using the RNeasy Mini Plus kit (Qiagen) according to the manufacturer’s instructions. Briefly, 5ml or 3ml RLT buffer containing 143 mM β-mercaptoethanol was added directly to the nasal tissue samples in gentleMACS M tubes (Miltenyi Biotec). Samples were homogenized using the gentleMACS dissociator (Miltenyi Biotec) and lysates were clarified twice by centrifugation at 3,000 ×*g* for 10 min. RNA was extracted using the animal tissue protocol and eluted in 50μl. RNA concentration and purity was assessed by spectrophotometry (*A*_260_/*A*_280_).

### RNA extraction and DNAse-treatment for ONT

Total RNA was extracted from cellular material using the RNeasy Mini Kit (Qiagen), using the manufacturer’s guidelines. The extracted RNA was treated with DNase using the Turbo DNA-free kit (Ambion). QC was carried out using the Qubit RNA High Sensitivity Assay (ThermoFisher Scientific) and Qubit 1 x dsDNA High Sensitivity Assay (ThermoFisher Scientific) on the Qubit 4 Fluorometer (ThermoFisher Scientific), High Sensitivity RNA and High Sensitivity D5000 Assays on the Tapestation 4200 System (Agilent Technologies) and NanoDrop™ 2000/2000 Spectrophotometer (ThermoFisher Scientific).

### ONT sequencing

DNAse-treated RNA was used as input for the PCR-cDNA Barcoding Kit (SQK-PCB111.24, ONT) according to the manufacturer’s guidelines, with minor modifications. Modifications include the increased input RNA amount to 500ng instead of the 200ng in the protocol, as well as using 18 cycles and 10 min extension times for the PCR step. Additionally, synthetic RNA (Sequin MixA) was added at 5% of the expected level of mRNA (5% total RNA) (41). PCR artifacts were removed using the prescribed method from ONT. The sequencing was carried out with R9.4.1 MinION flow cells on the ONT GridION for 72 hr.

### Data analysis

#### Basecalling

Original passing threshold fast5 (Qscore >= 10) output from the GridION was converted to pod5 format using pod5 tool to boost the efficiency of basecalling. *Dorado* (v0.5.3 or v0.9.6) was used to perform basecalling on the pod5 signals with the output format as ubam for quantification and poly(A) tail length estimation, respectively. “dna_r9.4.1_e8_sup@v3.6” was the selected model and “--estimate-poly-a” parameter was raised to estimate poly(A) tail length for each read in the sequence. “samtools (v1.16.1) fastq” command converted the ubam to fastq with ‘-T *’ parameter to keep all the tag information (42).

#### Restranding and reference mapping

Before reference mapping*, Restrander* (v1.0.1) (43) helped to remove artifact reads in our ONT long-read cDNA sequences, using the prebuild configuration for the PCB111 protocol. Reference alignment was performed by *minimap2* (v2.24) (44) with parameters as “-y -t 20 --eqx -k 12 -uf -ax splice --secondary=no”. The ferret reference genome used was downloaded from NCBI (Genome assembly ASM1176430v1.1, taxon as *Mustela putorius furo* (domestic ferret)). To keep the primary mapping reads only, “samtools view -F 2308” was run to filter for the mapped reads.

#### Quantification and isoform discovery

*Bambu* (v3.2.4) package in R was performed for multi-sample transcript discovery and quantification (45). Gene and transcript expressions were calculated for each sample. The extended annotation output from *Bambu* was used as input for *SQANTI3* (v5.1.2) (46). *Apptainer* (v1.2.3) was used to drive the *SQANTI3* container (47). “sqanti3_qc.py” and “sqanti3_filter ml” were executed step by step to first classify all isoforms and them using machine learning filter to remove artifacts. Then, the ‘rescue’ function was utilized to recover some transcripts in the artifact group. The quantification was carried out again using *Bambu* to generate the final count matrices and QC was carried out using *SQANTI3*.

*SQANTI3* isoform categories include Full Splice Match (FSM), Incomplete Splice Match (ISM), Novel In Catalog (NIC), Novel Not In Catalog (NNIC), Antisense, Genic Intron, Genic Genomic, and Intergenic. Each of these classes can be broadly defined as below:

FSM – all splice junctions match perfectly to the reference transcript

ISM – the splice junctions match partially to the reference transcript

NIC – novel isoform formed from a combination of known splice sites

NNC – novel isoform with at least a new splicing site

Genic Intron – transcript present within an intron

Genic Genomic – transcript with overlap in intronic and exonic regions

Intergenic – transcript present in an intergenic region

Novel genes and transcripts were visualized using *IGV* (v2.10.1) (48) and *IsoVis* (49). The novel ISGs and ISTs which were selected for further validation via PCR were determined by isolating a list of BambuGenes and BambuTxs which were significantly upregulated in the *in vitro* comparison between the IFN-treated vs mock-control groups (padj < 0.05, see ‘**differential expression and polyadenylation analyses’**). Then, the mean expression level based on the transcript counts determined by *Bambu* between sequencing replicates were defined and the ISGs and ISTs were ranked based on this average statistic. For the ISGs, the low, mid, high genes were selected based on their ranking positioning. For the ISTs, the transcripts were selected based on whether the transcript was from an annotated gene and ranked based on the positions within this subset.

#### PolyA tail length estimation

The estimated poly(A) tail lengths of each read were stored in the ubam files from the basecalling step. A combination of “samtools view” and bash commands were implemented to extract the poly(A) tail lengths and sequence lengths into a text format via “samtools view my.bam | awk ‘/pt:i/{print $1,length($10),$NF-1}’ | sed ‘s/pt:i://g’”.

#### Differential expression and polyadenylation analyses

Counts for duplicate transcripts were first merged in the counts matrices for transcripts. *DEseq2* (v1.38.3) (50) package in *R* (v4.2.1) was utilized to test differential expression between different conditions, where a padj threshold of 0.05 was used. *ggplot*, *pheatmap, and ComplexHeatmap* packages in *R* were used to produce the volcano plots and heatmap plots.

The differential polyadenylation analysis aims to identify variants in poly(A) tail lengths across different sample conditions. By comparing two sets of paired conditions in both *in vivo* and *in vitro* samples, the analysis elucidated changes in polyadenylation patterns. From the basecalled data, *IsoQuant* (v3.3.1) (51) was employed to assign cDNA reads to genes. After cross-matching poly(A) tail length information and gene assignment, we selected genes with more than 100 reads assigned and at least 10 reads assigned to a single sequencing replicate included in the comparison. The *R* package *lmerTest* (v3.1.3) (52) conducted a linear mixed-effects regression (lmer), where the log-transformed poly(A) length for all reads mapped to one gene served as the response variable, while the type of infection or treatment as the fixed effects and sample batch as the random effects. The per-gene P-values were generated and adjusted using the Benjamini-Hochberg (BH) method with the ‘p.adjust’ function in *R*. The differences between mean of poly(A) tail length from each condition on each gene were calculated using *R*. Volcano plots were constructed using the *R* package *ggplot2* (v3.5.1) by setting the p.adjust value cut off ≤ 0.05 as significant.

#### Gene Set Enrichment and KEGG Pathway Enrichment Analyses

GSEA analysis for polyadenylation was implemented with the *ClusterProfiler* package in *R* (53). The input data was the lmerTest gene list ordered by difference of the average poly(A) tail for a condition (‘diff’) for each comparison in differential polyadenylation analysis. The KEGG database for the *Mustela putorius furo* (domestic ferret) (https://www.kegg.jp/kegg-bin/show_organism?menu_type=pathway_maps&org=mpuf) was implemented for the enrichment analysis. For the KEGG pathway analysis, significant genes discovered in the differential polyadenylation analyses were queried in the enrichment analysis with background gene lists containing only genes which were involved in the differential analysis. Significant genes were separated into subgroups by upregulation and downregulation in different comparison groups.

#### Protein structure analysis

The sequences of hypothetical trans-spliced transcripts were analyzed for potential open reading frames (ORF) via the NCBI *ORFfinder* tool. The most robust protein sequences based on existing reference *IFITM* transcripts were selected for protein structure predictions via the *trRosetta* tool (54).

#### RT-qPCR data analysis and gene alignments

Graphs and statistical analysis (as indicated in the figure legends) was generated using R-Studio. Statistical analysis was performed by utilizing a mixed effects model (*nlme* package in *R*). Protein sequence alignments were performed using *MUSCLE* alignment function on *Geneious Prime®* (v2025.0.2). The sequences used for the protein alignment are as follows (of note the ORF sequence from the ferret transcripts were first translated *in silico*):

| Gene | Ferret | European Mink | Humans | Mouse |
| --- | --- | --- | --- | --- |
| BST2 | XM_045081929.1 | XP_059016426.1 | NP_004326.1 | NP_932763.1 |
| ISG15 | XM_004781282.3 | XP_058994206.1 | NP_005092.1 | NP_056598.2 |
| Viperin | XM_004745742.3 | XP_058988730.1 | Isoform 1:<br>NP_542388.2, Isoform 2:<br>NP_001397631.1 | NP_067359.2 |
| MOV10 | Isoform 1:<br>XM_045081891.1,<br>Isoform 2:<br>XM_004769794.3 | XP_058993685.1 | NP_001123551.1 | NP_032645.2 |
| OAS2 | XM_004753413.2, region-3456-6750 | XP_058995097.1 | Isoform 1:<br>NP_058197.2, Isoform 2:<br>NP_002526.2 | Isoform 1: NP_660262.2,<br>Isoform 2:<br>NP_001334377.1 |
| TRIM22 | XM_004778994.3 | XP_059038208.1 | Isoform 1:<br>NP_006065.2, Isoform 2:<br>NP_001186502.1 | N/A |

#### Data availability

All raw fastq sequencing datasets with poly(A) tail length tags are available on NCBI SRA at Project PRJNA1290003 and scripts are available on Github - https://github.com/abcdtree/ferret-rna-ont-paper.

