## Supplementary Figures for "Characterizing the transcriptomic response to interferon and infection in European Domestic Ferret respiratory tissues using long-read RNA sequencing"

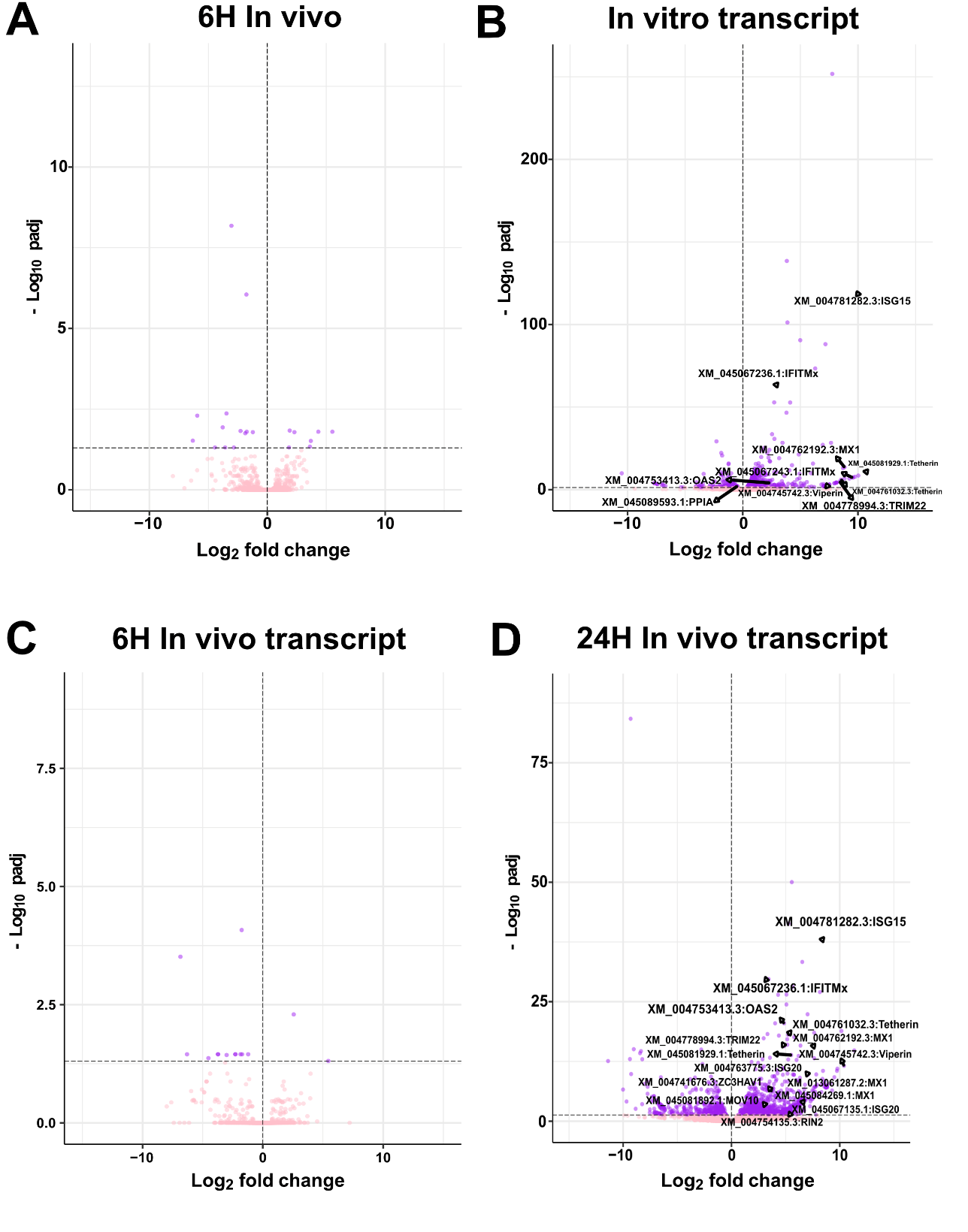

**Figure S1. Upregulated human ISG/HRF-orthologous genes and transcripts in ferrets comparing IFNα-treated vs mock-control ferret FRL cells (*in vitro*) and nasal turbinate samples from IAV-infected vs mock-infected control ferrets (*in vivo*) at 6 and 24 hpi. A)** Differential expression results between 6 hpi IAV-infected vs mock-infected control data (*in vivo*) at the gene level. **(B-D)** Differential transcript expression results between **B)** IFNα-treated vs mock-control FRL cells (*in vitro*), and **C)** 6 hpi (*in vivo*) and **D)** 24 hpi IAV-infected ferret vs mock-infected control data (*in vivo*). X-axis represents log_2_ fold changs and the Y-axis shows the -log_10_ *p*-adjusted value (padj), with a threshold of padj < 0.05, shown by the dotted horizontal line.

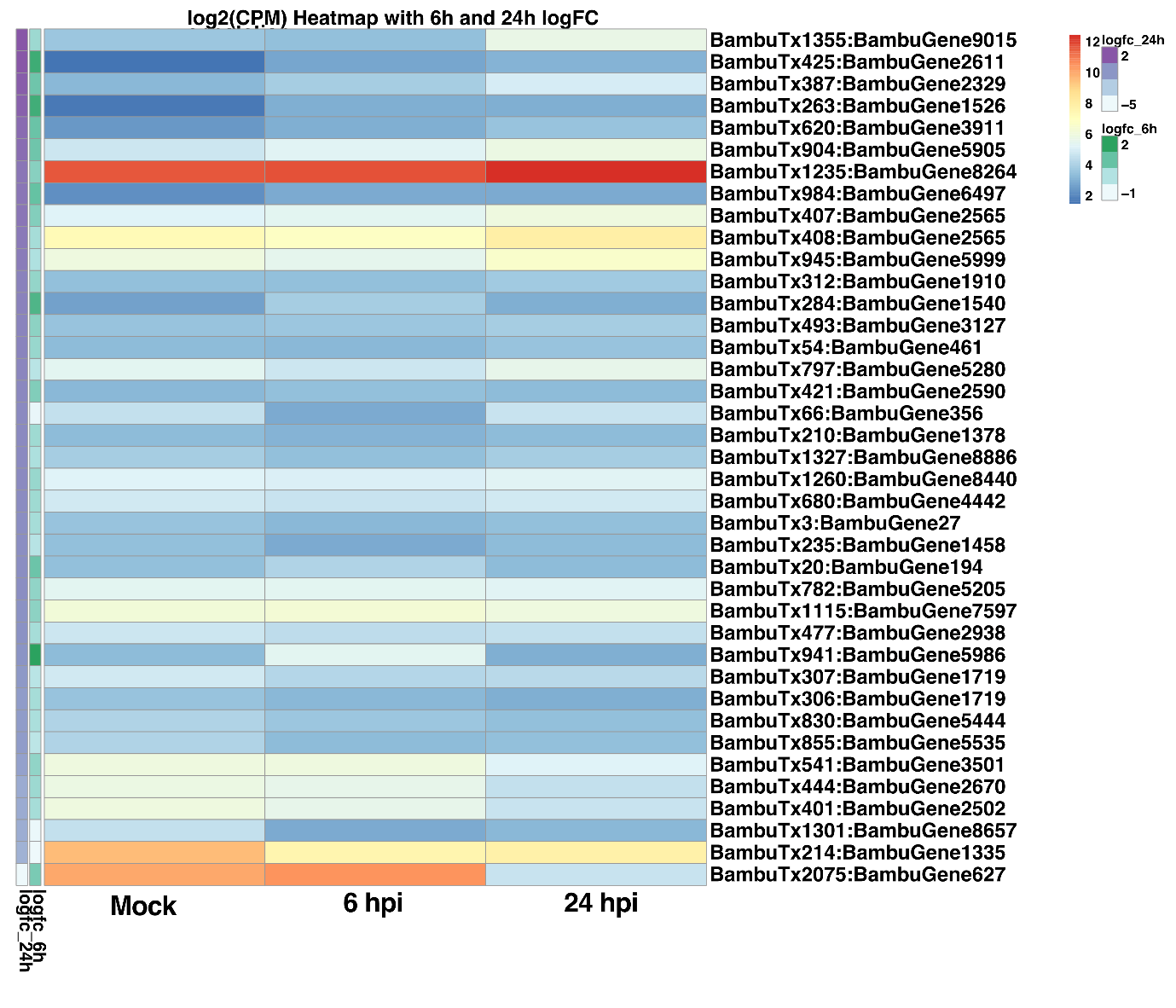

**Figure S2.** **Heatmap of log₂-transformed gene expression values (CPM) across three *in vivo* conditions: mock-treated, 6 and 24 hr post-infection (hpi).** Each row represents a gene, and columns represent the conditions. Expression patterns are annotated with two side colour bars representing logFC values at 6 and 24 hpi, respectively. Increase in purple intensity indicates higher logFC values at 24 hpi and increases in green intensity indicates higher logFC values at 6 hpi. Genes are ordered by decreasing logFC at 24 hpi. No row or column clustering was applied to preserve condition and ranking order.

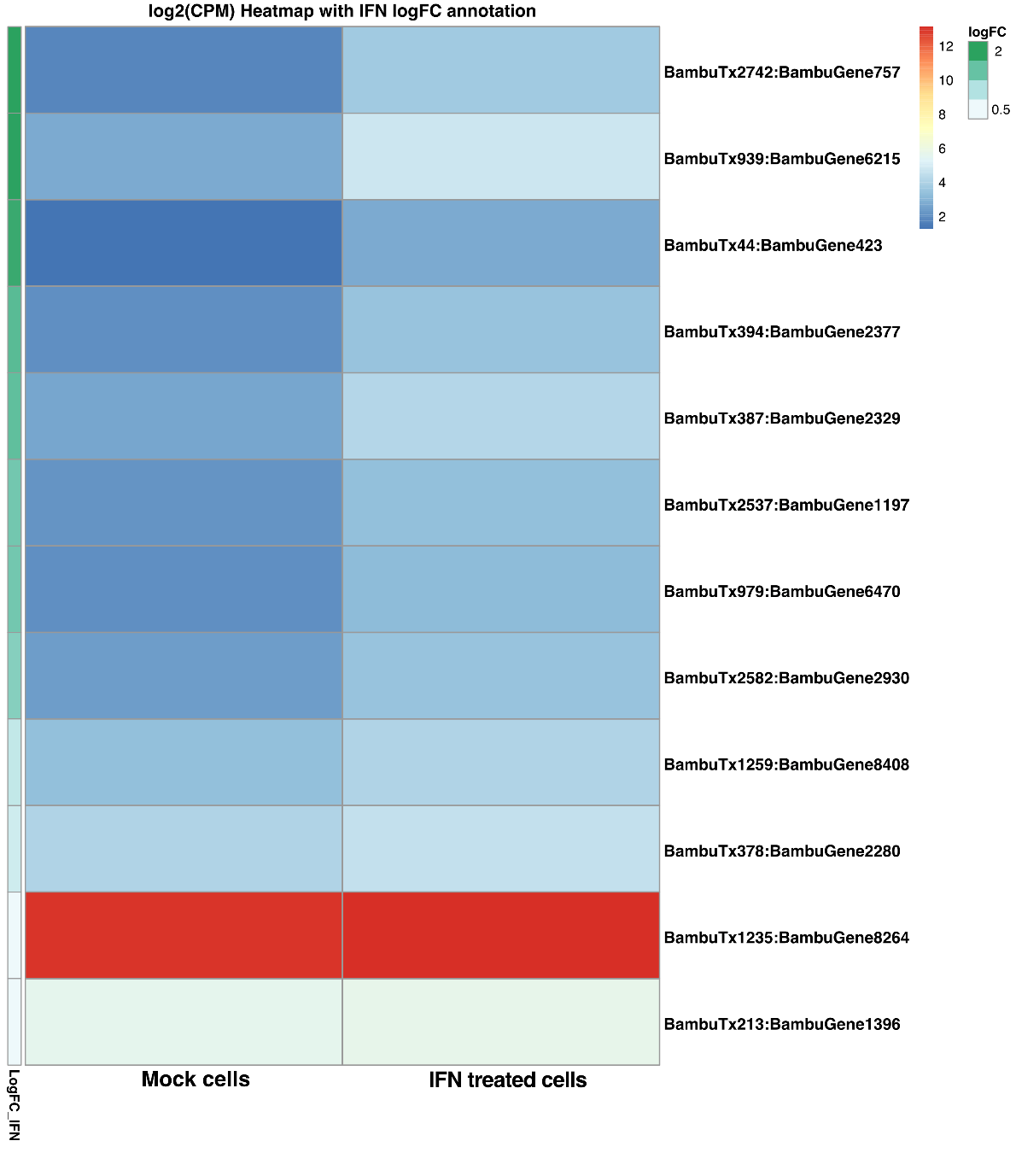

**Figure S3.** **Heatmap showing the log₂-transformed expression levels of selected genes in mock-treated and IFN-treated FRL cells from *in vitro* experiments.** Expression values are based on CPM (counts per million) and are displayed for each gene across the two conditions. Genes are ordered by descending log fold change (logFC) in response to IFN treatment. The side colour bar represents logFC values, where green indicates strong upregulation following IFN treatment. Rows are not clustered to preserve the order of logFC.

**A) Primer binding sites for IFITM transcript 1, 2 and 3**

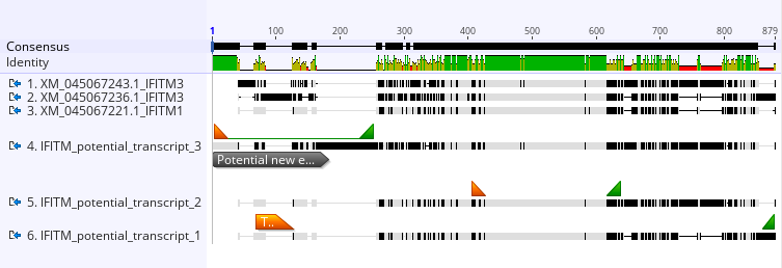

**B) Primer binding site for IFITM transcript 2**

Forward	Reverse

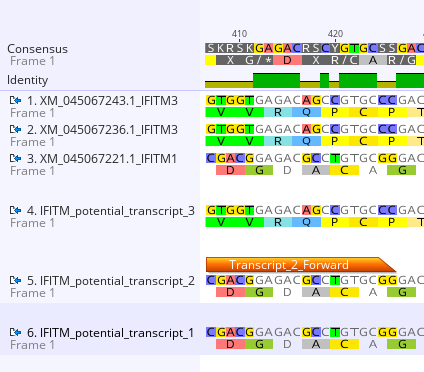

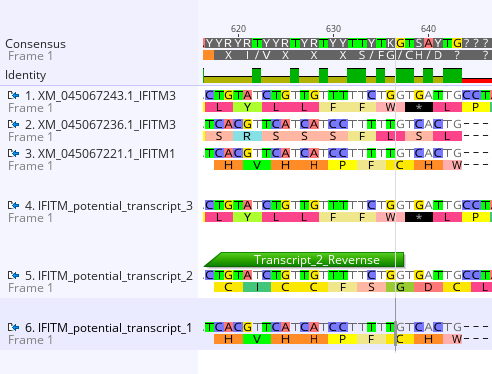

**Figure S4.** **Location of primer binding sites for novel trans-spliced IFITM transcripts. A)** The forward (orange) and reverse (green) primers for transcripts 1 and 3 bind to regions unique to the respective novel trans-spliced transcripts. **B)** For transcript 2, although the individual forward (orange) and reverse (green) primers are not unique on their own, their specific combination produces a unique amplification product for transcript 2.
